# Your Emotions, My Brain: Generalizable Neural Signatures of Emotional Memory Reactivation During Sleep

**DOI:** 10.1101/2025.08.11.669349

**Authors:** Mahmoud E. A. Abdellahi, Tia Tsimpanouli, Penelope A. Lewis

## Abstract

Reactivation in sleep alters the structure of memories and can potentially be used to restructure upsetting representations. Reactivation can be triggered with auditory cues and then detected using machine learning and electroencephalography (EEG), but can we also detect the emotionality of reactivated memories? We examined this by presenting auditory cues that had been associated with negative or neutral stimuli in wake during subsequent NREM sleep and training a classifier to detect the emotionality of subsequent EEG responses. We were able to detect the reinstatement of emotionality 0.4-0.6 seconds after cue presentation. Importantly, we used a between-participant machine learning pipeline to identify shared neural signatures of emotionality across individuals without fine-tuning the model on testing participants. This approach eliminates the need for individualized wake localizer sessions, establishing a methodologically efficient framework for investigating emotional processing during sleep. Detection of emotional reactivation in sleep will help us to understand how such reactivation impacts upon the emotionality of the memories in the long term, potentially facilitating development of treatments for PTSD and depression through memory restructuring in sleep.

## Introduction

Maladaptive emotional memories contribute to psychiatric symptoms in depression and PTSD (Germain, 2013; Walker & van der Helm, 2009). Intentional reactivation of emotional memories during sleep using Targeted Memory Reactivation (TMR) has been shown to improve therapy outcomes (van der Heijden et al., 2024) and reduce emotionality of upsetting memories (Greco et al., 2024; Hutchison et al., 2021a; Wassing et al., 2019; Xia et al., 2023, 2024). A tool for the detection of emotional reactivation in sleep would greatly increase our ability to study the impact of such reactivation upon emotional processing, and to harness such interventions for clinical benefit. Both multivariate EEG decoding (Abdellahi et al., 2023a; Cairney et al., 2018; Schreiner et al., 2018) and fMRI-based activation patterns (Rasch et al., 2007; Shanahan et al., 2018) have established that TMR can successfully trigger content-specific memory reactivation during both slow-wave and REM sleep. However, no study has yet attempted to detect the emotionality or affective charge of such reactivation events. In this paper, we set out to develop a classifier which can detect the emotionality (negative or neutral) of memory reactivation in NREM sleep.

In order to detect memory reactivation during sleep, it is necessary to train a classifier during wakeful experiencing of the memory in question. This typically requires a repetitive and time-consuming localiser task which not only limits experimental throughput but also constrains our understanding of universal principles of sleep-dependent memory consolidation, as findings remain tied to patterns of individual participants rather than revealing generalisable neural mechanisms. In the case of emotional memory, to which many responses habituate with repeated experiences of the emotional stimulus, such localisers can be problematic. An alternative would be to train a classifier that generalises across participants such that, once trained, it could be applied on new emotional TMR studies without the need for a localiser. While no study has, as yet, trained a generalisable classifier to identify emotionality of memories in sleep, prior work has successfully trained classifiers to identify emotions in wake (Ding et al., 2025; Jiménez-Guarneros & Fuentes-Pineda, 2025; Shi et al., 2024), suggesting that such processing has a common enough physiological basis to lend itself to such work.

To achieve this, we investigated whether the emotionality of images used to cue finger movements in a serial reaction time task (SRTT) (Cousins et al., 2014, 2016), a task which is known to reactivate after TMR cues (Abdellahi et al., 2023a, 2023b; Belal et al., 2018), can be detected using EEG classifiers when these memories are reactivated during NREM sleep. Participants learned two SRTT sequences—one paired with negative emotional stimuli (fearful faces and aversive objects) and the other with neutral stimuli (neutral faces and objects). During subsequent NREM sleep, we presented auditory cues associated with both sequences and analysed the resulting EEG activity.

Theta power at 4-8 Hz has been observed to increase following successful memory cues during sleep (Lehmann et al., 2016), and theta oscillations appear to coordinate memory reactivation events during NREM sleep (Schreiner et al., 2018). Theta oscillations may therefore be especially relevant for emotional memory processing (Legendre et al., 2022; Lehmann et al., 2016). We therefore used theta power as the feature of interest and developed a machine learning approach to classify the emotional context of reactivated memories, employing a between-participant generalization method to identify shared neural signatures across individuals.

## Results

We developed a classification approach using theta band (4-8 Hz) power features to discriminate between the reactivation of negative and neutral SRTT sequences during NREM sleep following TMR. Our machine learning pipeline incorporated instantaneous theta power features extracted using Hilbert transformation and assessed using a leave-one-participant-out cross-validation approach with linear discriminant analysis (LDA) classifiers. Unlike previous work that typically analysed classification within individual participants(Belal et al., 2018; Cairney et al., 2018), our approach addressed the generalizability of emotional memory signatures across participants.

We found that the emotional context associated with a TMR cue can be successfully classified during NREM sleep following the cue (Fig. 2). Classification performance measured by area under the receiver operating characteristic curve (AUC) revealed significant classification of negative versus neutral context (p = 0.0038), with the strongest classification occurring between 0.4-0.6 seconds after cue presentation. This timing is consistent with previous work examining neural responses to memory cues during sleep, showing an increase in theta activity during a similar timing is associated with improved memory performance (Schreiner et al., 2015).

**Figure 1.**
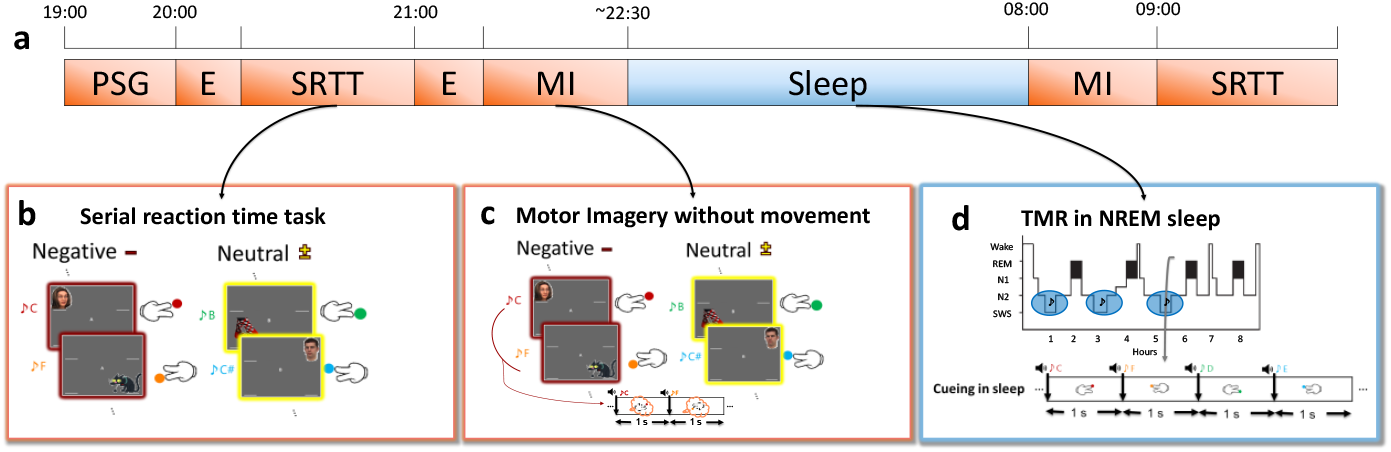
Experimental design. (**a**) Timeline of the experiment showing the sequence of tasks. Participants were fitted for polysomnography (PSG) recording before completing an engagement task (E), the serial reaction time task (SRTT), another engagement task, and a motor imagery (MI) task. Subsequently, they spent overnight sleep in the lab with targeted memory reactivation carried out. In the following morning, participants performed the motor imagery task and SRTT again. (**b**) The SRTT contained two distinct sequences associated with different emotional contexts. The negative sequence (red frame) included faces (top corners) or objects (bottom corners), faces were a fearful female face and a sad male face, objects were a dirty toilet and a severed arm. The actual images were not used for viewers’ convenience. The neutral sequence (yellow frame) contained the same individuals with neutral expressions and neutral objects (wooden chair and socks). Each image appeared in a specific corner of the screen accompanied by a specific tone, requiring participants to press the button corresponding the image corner. (**c**) In the motor imagery task, participants viewed the same sequences as in the SRTT but were instructed to only imagine pressing the buttons without actual movement. (**d**) During NREM sleep, tones associated with both sequences were presented TMR was performed after at least 20 minutes of stable NREM sleep, with stimulation resumed if arousals or sleep stage changes occurred.

**Figure 2.**
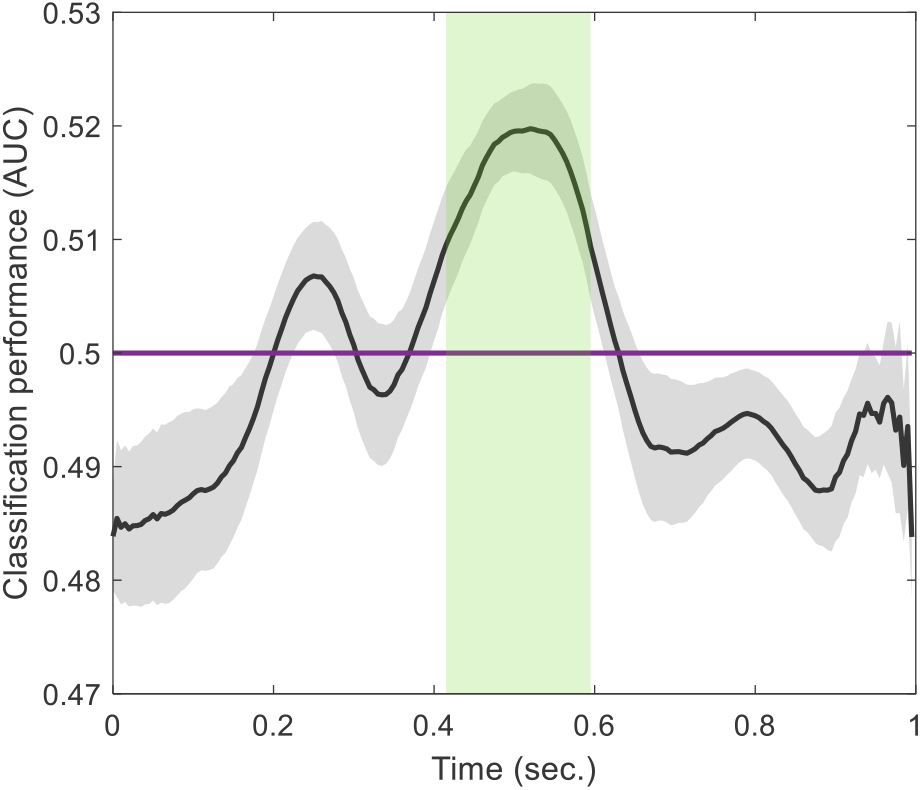
Classification performance for negative versus neutral emotional contexts during NREM sleep following TMR. The plot shows the area under the receiver operating characteristic curve (AUC) over time after TMR cue presentation, with the black line representing the mean classification performance across participants and the grey shaded area indicating the standard error. Significant classification (p = 0.0038) occurred between 0.4-0.6 seconds after cue presentation (highlighted in green). The purple horizontal line at 0.5 represents chance-level classification. This time window corresponds to the period when emotional context is most strongly encoded in theta power features, suggesting a consistent temporal profile of emotional memory reactivation across participants.

Importantly, the successful between-participant generalization demonstrates that the neural signatures of emotional memory reactivation share common features across individuals, providing evidence for a generalizable mechanism of emotional memory processing during sleep. These findings suggest that the emotional valence associated with procedural memories is preserved during sleep-based reactivation and can be detected using EEG classifiers trained on theta power features.

## Discussion

Our findings demonstrate that the emotional context associated with procedural memories can be detected during sleep-based reactivation with EEG classifiers. By employing theta power features and a leave-one-participant-out cross-validation approach, we were able to successfully distinguish between the reactivation of memories with negative versus neutral emotional charge during NREM sleep following TMR. These results provide new insights into how emotional information is processed during sleep and suggest common neural mechanisms for emotional processing in sleep across individuals.

The creation of a machine learning pipeline which can detect emotional signatures using a generalized classification model trained across participants and knowing the precise timing of that reactivation after TMR cues represents a methodological advancement in the study of sleep-based memory processing. The current pipeline does not rely on fine-tuning which involves adapting to samples from the testing participant, this makes it a zero-shot model meaning that it can be directly applied to new unseen participants. Previous work has primarily employed within-participant classification approaches (Abdellahi et al., 2023a; Belal et al., 2018; Cairney et al., 2018; Schreiner et al., 2023) which do not address whether memory reactivation patterns share common features across individuals. Our findings suggest that emotional memory processing during NREM sleep involves generalizable neural mechanisms that can be identified using machine learning techniques, potentially opening new avenues for investigating sleep-dependent memory consolidation.

Importantly, each sequence contained 12 different sounds, and the association between sequences and emotional content (negative or neutral) was randomized across participants. Thus, the successful classification across participants strongly suggests that our model captured neural signatures specific to emotional charge rather than superficial stimulus properties.

The observed classification effect occurring 0.4-0.6 seconds after cue presentation, suggests temporal consistency of reactivation between trials and individuals, with a critical period for the unfolding of memory reactivation processes following external cues. Future research should investigate how this temporal pattern relates to other oscillatory phenomena during NREM sleep, particularly slow oscillations and sleep spindles, which are known to play crucial roles in memory consolidation (Helfrich et al., 2019; Schreiner et al., 2021, 2023; Staresina et al., 2015). It is suggested that memory reactivation is nested within the hierarchical organization of these oscillations (Klinzing et al., 2019), and exploring how emotional memory processing interacts with these rhythms could provide further insights into the mechanisms of sleep-dependent consolidation.

Future work could enhance classification performance by implementing methods to align neural data between individuals. Analogues to fMRI techniques such as hyperalignment (Haxby et al., 2011) EEG-based techniques specifically from the domain of contrastive learning could be valuable for this purpose (Shen et al., 2024). Attention-based models could further help pinpoint the specific neural representations that are most relevant for emotional memory processing during sleep (Vaswani et al., 2017), allowing us to identify which features are shared across participants versus those that differ. By mapping the commonalities and differences in how individuals process emotional memories, such approaches could not only improve classification performance but also contribute to our understanding of the neural mechanisms underlying emotional memory consolidation. These alignment methods could also reveal which aspects of theta oscillations are consistently involved in emotional memory processing across individuals, potentially identifying biomarkers for successful memory reactivation and consolidation.

Given the well-established relationship between REM sleep and emotional memory processing (Greco et al., 2024; Groch et al., 2013, 2015; Hutchison et al., 2021b; Wiesner et al., 2015), an important next step will be to investigate whether similar classification approaches can detect emotional memory signatures during REM sleep. Comparing the neural signatures of emotional memory reactivation across different sleep stages could provide valuable insights into the complementary roles of NREM and REM sleep in emotional memory processing.

The consolidation of maladaptive emotional memories plays a significant role in conditions like depression and PTSD (Germain, 2013; Walker & van der Helm, 2009). Recent studies have indicated that TMR can actually improve the outcomes of trauma therapy (van der Heijden et al., 2024), while other work has shown that TMR can be used to make emotional memories less upsetting (Greco et al., 2024; Hutchison et al., 2021a; Schwartz et al., 2022; Wassing et al., 2019), and to ‘update’ negative memories by strengthening the association with more positive information (Xia et al., 2023, 2024). Overall, the literature suggests a real potential for TMR as a therapeutic tool (Cabrera et al., 2024). Developing effective ways to specifically detect reactivation of the emotional components of these memories is a critical next step in terms of understanding how such reactivation might be used to impact on subsequent mood and potentially to alleviate psychiatric symptoms.

In conclusion, we have demonstrated that the emotional charge associated with negative memories is preserved during sleep-based reactivation and can be detected using EEG classifiers. Our classifier is the first to achieve between-participant classification in sleep classifying emotional content and therefore provides a useful tool for further studies of emotional processing in sleep. A better understanding of how emotional processing can be intentionally triggered through external manipulations could facilitate the development of TMR based treatments for disorders of emotional memory, such as PTSD and depression.

## Methods

### Participants

EEG and behavioural data were collected from 19 healthy women (mean age = 23.58 years, SD 4.5). As evidence suggests that women and men use different strategies to process emotional information and show differences in which brain areas get activated (Bianchin & Angrilli, 2012). All participants had normal hearing, normal or corrected-to-normal vision, reported a normal sleeping pattern for at least 4 weeks before participating, and abstained from caffeine and alcohol for at least 24 hours prior to the experiment. Eighteen participants were right-handed and one was left-handed. No participants reported having a history of sleep, motor, neurological, or psychiatric disorders, or taking any psychologically active medications. All participants provided written informed consent.

### Experimental design

Upon arrival at approximately 7 PM, participants completed a series of questionnaires to assess their sleep quality and habits, as well as their rumination, depression, anxiety, and stress levels. These included the Morningness-Eveningness Questionnaire (MEQ), a Sleep Diary, the 22-item Ruminative Responses Scale (RRS), and the 21-item Depression Anxiety and Stress Scale (DASS). Participants were then fitted for polysomnography (PSG) recording and began the experimental tasks around 8 PM.

The SRTT was adapted from (Cousins et al., 2014, 2016) and contained interleaved blocks of two 12-item sequences (A: 1 2 1 4 2 3 4 1 3 2 4 3 and B: 2 4 3 2 3 1 4 2 3 1 4 1) that were matched for learning difficulty. Each block contained 3 repetitions of a sequence, and no more than two blocks of the same sequence were presented consecutively. Blocks were separated by a 15-second gap during which performance feedback was displayed. Participants performed 24 blocks of each sequence and an additional 4 blocks with random sequences.

Each sequence was associated with either negative or neutral visual cues, accompanied by 200ms long high- or low-pitched pure tones, all counterbalanced across participants. Visual cues included four objects from the International Affective Pictures System (IAPS)(Bradley & Lang, 2017) or the internet, and four faces from the Radboud Faces Database (Langner et al., 2010). The neutral objects were a wooden chair and a pair of legs wearing socks, while the corresponding negative objects were a dirty toilet and a severed arm. The neutral faces were one male and one female with neutral expressions, while the negative faces were the same individuals with the female showing a fearful expression and the male showing a sad expression.

For each trial, a visual cue appeared in one of the four corners of the screen accompanied by a tone. Semantically related visual cues appeared in the same position for each sequence (1– top left corner=female face, 2–bottom left corner=body, 3–top right corner=male face, 4– bottom right corner=sitting object). Participants pressed the corresponding key (left shift, left Ctrl, up arrow, or down arrow) as quickly as possible while minimizing errors. Visual cues disappeared only after a correct response and were followed by an 880ms inter-trial interval.

Prior to the SRTT and motor imagery tasks, participants completed a short engagement task designed to enhance the emotional salience of the stimuli. In this task, participants received appropriate instructions (e.g., mimicking facial expressions, imagining themselves in a situation using the presented object) before each stimulus was displayed in full screen for 5 seconds.

Following the SRTT, participants performed a motor imagery (IMG) task in which they viewed the same sequences of images but only imagined pressing the corresponding keys without actual movement. This task consisted of 48 interleaved blocks (24 from each sequence) presented in the same order as in the SRTT. Cues were presented for 270ms followed by an 880ms inter-trial interval, and no feedback was provided between blocks.

After the evening experimental tasks, participants were allowed to read in bed prior to sleep. The next morning, participants were awoken after 7-8 hours of sleep and allowed 20 minutes to overcome sleep inertia before completing the morning IMG task followed by the SRTT.

### PSG data acquisition and analysis

Silver-silver chloride electrodes were attached to the scalp and face of the participants according to the 10-20 system. In total, 23 electrodes were placed, with 16 electrodes on standard scalp locations (F3, F4, C3, C4, Cz, C5, C6, CP3, CP4, CP5, CP6, P7, P8, Pz, O1, and O2), referenced to the contralateral mastoid electrode. Additional electrodes were placed next to the eyes, on the chin, and at the forehead (ground). Connection impedance was <5KOhms and the sampling rate was 200Hz. PSG activity was recorded using an Embla N7000 PSG amplifier with RemLogic software. All PSG recordings were scored manually by two experimenters who were blind to when sounds were cued, according to “The AASM Manual for the Scoring of Sleep and Associated Events”.

### TMR delivery during NREM sleep

Brown noise was played throughout the night to minimize any sleep disturbances caused by external noises. Replay of sequence tones began after at least 20 minutes of stable NREM sleep. For all participants, tones from both sequences (negative and neutral) were presented with the same duration of cue presentation and inter-trial interval as in the imagery task.

Tones of the two sequences were replayed in alternating blocks. For six participants, each block contained one repetition of a sequence, while for thirteen participants it contained five repetitions. The interval between two sequences was at least 20 seconds long. If an arousal occurred during replay or a sleep stage change was detected, replay was paused immediately and resumed at least one minute later or when participants returned to stable NREM sleep.

### Detection of emotional context with machine learning

We developed a classification approach to detect the emotional context (negative versus neutral) of reactivated memories during NREM sleep after TMR. Following previous successful work of linear classification for reactivation detection (Abdellahi et al., 2023b, 2023a), we used Linear discriminant analysis (LDA) classifiers trained and tested on each time point after TMR cues. EEG signals were band-pass filtered in the theta range (4-8 Hz) and Hilbert transformation was applied to extract instantaneous power values. For each trial from every participant, we calculated that instantaneous theta power resulting in the same dimensions as the original data (trials x channels x timepoints). These power features were used as inputs for classification. One classification model was trained at a time point and applied to testing set at that same time point.

We employed a leave-one-participant-out cross-validation approach to assess the generalizability of emotional memory signatures across individuals. LDA classifiers were trained on data from n-1 participants (16 participants) and tested on the left-out participant, with this process repeated for all 17 participants. This approach differs from within-participant classification methods used in previous studies (Belal et al., 2018; Cairney et al., 2018), allowing us to examine whether neural signatures of emotional memory reactivation share common features across individuals.

To evaluate classification performance across time, we applied the LDA classifier at each time point after the TMR cue (King & Dehaene, 2014). This time-resolved classification allowed us to identify when emotional context was most strongly encoded in the theta power features. Classification performance was quantified using the area under the receiver operating characteristic curve (AUC), with statistical significance determined using cluster-based permutation testing to correct for multiple comparisons (Oostenveld et al., 2011).

## Notes

### Competing Interest Statement

The authors have declared no competing interest.

### Funding

- Wellcome Trust: 227012/Z/23/Z

## References

Abdellahi, M. E. A., Koopman, A. C. M., Treder, M. S., & Lewis, P. A. (2023a). Targeted memory reactivation in human REM sleep elicits detectable reactivation. ELife, 12. 10.7554/ELIFE.84324

Abdellahi, M. E. A., Koopman, A. C. M., Treder, M. S., & Lewis, P. A. (2023b). Targeting targeted memory reactivation: characteristics of cued reactivation in sleep. NeuroImage, 266, 2021.12.09.471945. https://www.sciencedirect.com/science/article/pii/S1053811922009417?via%3Dihub

Belal, S., Cousins, J., El-deredy, W., Parkes, L., Schneider, J., Tsujimura, H., Zoumpoulaki, A., Perapoch, M., Santamaria, L., & Lewis, P. (2018). Identification of memory reactivation during sleep by EEG classification. NeuroImage, 176(December 2017), 203–214. 10.1016/j.neuroimage.2018.04.029

Bianchin, M., & Angrilli, A. (2012). Gender differences in emotional responses: A psychophysiological study. Physiology & Behavior, 105(4), 925–932. 10.1016/J.PHYSBEH.2011.10.031

Bradley, M. M., & Lang, P. J. (2017). International Affective Picture System. Encyclopedia of Personality and Individual Differences, 1–4. 10.1007/978-3-319-28099-8_42-1

Cabrera, Y., Koymans, K. J., Poe, G. R., Kessels, H. W., Van Someren, E. J. W., & Wassing, R. (2024). Overnight neuronal plasticity and adaptation to emotional distress. Nature Reviews Neuroscience, 25(4), 253–271. https://doi.org/10.1038/S41583-024-00799-W;SUBJMETA=378,631,692,700;KWRD=HEALTH+CARE,NEUROSCIENCE

Cairney, S. A., Guttesen, A.á.V., El Marj, N., & Staresina, B. P. (2018). Memory Consolidation Is Linked to Spindle-Mediated Information Processing during Sleep. Current Biology, 28(6), 948-954.e4. 10.1016/j.cub.2018.01.087

Cousins, J. N., El-Deredy, W., Parkes, L. M., Hennies, N., & Lewis, P. A. (2014). Cued memory reactivation during slow-wave sleep promotes explicit knowledge of a motor sequence. Journal of Neuroscience, 34(48), 15870–15876. 10.1523/JNEUROSCI.1011-14.2014

Cousins, J. N., El-Deredy, W., Parkes, L. M., Hennies, N., & Lewis, P. A. (2016). Cued Reactivation of Motor Learning during Sleep Leads to Overnight Changes in Functional Brain Activity and Connectivity. PLoS Biology, 14(5), e1002451. 10.1371/journal.pbio.1002451

Ding, Y., Tong, C., Zhang, S., Jiang, M., Li, Y., Lim, K. J. L., & Guan, C. (2025). EmT: A Novel Transformer for Generalized Cross-Subject EEG Emotion Recognition. IEEE Transactions on Neural Networks and Learning Systems, 36(6), 10381–10393. 10.1109/TNNLS.2025.3552603,

Germain, A. (2013). Sleep disturbances as the hallmark of PTSD: Where are we now? American Journal of Psychiatry, 170(4), 372–382. 10.1176/APPI.AJP.2012.12040432,

Greco, V., Foldes, T. A., Harrison, N. A., Murphy, K., Wawrzuta, M., Abdellahi, M. E. A., Lewis, P. A., & Penelope, L. A. (2024). Disarming emotional memories using Targeted Memory Reactivation during Rapid Eye Movement sleep. BioRxiv, 2024.09.25.614960. 10.1101/2024.09.25.614960

Groch, S., Wilhelm, I., Diekelmann, S., & Born, J. (2013). The role of REM sleep in the processing of emotional memories: Evidence from behavior and event-related potentials. Neurobiology of Learning and Memory, 99, 1–9. 10.1016/j.nlm.2012.10.006

Groch, S., Zinke, K., Wilhelm, I., & Born, J. (2015). Dissociating the contributions of slow-wave sleep and rapid eye movement sleep to emotional item and source memory. Neurobiology of Learning and Memory, 122, 122–130. 10.1016/j.nlm.2014.08.013

Haxby, J. V., Guntupalli, J. S., Connolly, A. C., Halchenko, Y. O., Conroy, B. R., Gobbini, M. I., Hanke, M., & Ramadge, P. J. (2011). A common, high-dimensional model of the representational space in human ventral temporal cortex. Neuron, 72(2), 404–416. 10.1016/j.neuron.2011.08.026

Helfrich, R. F., Lendner, J. D., Mander, B. A., Guillen, H., Paff, M., Mnatsakanyan, L., Vadera, S., Walker, M. P., Lin, J. J., & Knight, R. T. (2019). Bidirectional prefrontal-hippocampal dynamics organize information transfer during sleep in humans. Nature Communications 2019 10:1, 10(1), 1–16. 10.1038/s41467-019-11444-x

Hutchison, I. C., Pezzoli, S., Tsimpanouli, M. E., Abdellahi, M. E. A., & Lewis, P. A. (2021a). Targeted memory reactivation in REM but not SWS selectively reduces arousal responses. Communications Biology 2021 4:1, 4(1), 1–6. 10.1038/s42003-021-01854-3

Hutchison, I. C., Pezzoli, S., Tsimpanouli, M. E., Abdellahi, M. E. A., & Lewis, P. A. (2021b). Targeted memory reactivation in REM but not SWS selectively reduces arousal responses. Communications Biology, 4(1). 10.1038/S42003-021-01854-3,

Jiménez-Guarneros, M., & Fuentes-Pineda, G. (2025). Multi-modal supervised domain adaptation with a multi-level alignment strategy and consistent decision boundaries for cross-subject emotion recognition from EEG and eye movement signals. Knowledge-Based Systems, 315, 113238. 10.1016/J.KNOSYS.2025.113238

King, J. R., & Dehaene, S. (2014). Characterizing the dynamics of mental representations: The temporal generalization method. Trends in Cognitive Sciences, 18(4), 203–210. 10.1016/J.TICS.2014.01.002/ASSET/34F4EF3B-CEA0-40D5-9749-9C25F22EF6FA/MAIN.ASSETS/GR1B1.SML

Klinzing, J. G., Niethard, N., & Born, J. (2019). Mechanisms of systems memory consolidation during sleep. In Nature Neuroscience. 10.1038/s41593-019-0467-3

Langner, O., Dotsch, R., Bijlstra, G., Wigboldus, D. H. J., Hawk, S. T., & van Knippenberg, A. (2010). Presentation and validation of the radboud faces database. Cognition and Emotion, 24(8), 1377–1388. 10.1080/02699930903485076;PAGE:STRING:ARTICLE/CHAPTER

Legendre, G., Bayer, L., Seeck, M., Spinelli, L., Schwartz, S., & Sterpenich, V. (2022). Reinstatement of emotional associations during human sleep: an intracranial EEG study. BioRxiv, 2022.06.24.497499. 10.1101/2022.06.24.497499

Lehmann, M., Schreiner, T., Seifritz, E., & Rasch, B. (2016). Emotional arousal modulates oscillatory correlates of targeted memory reactivation during NREM, but not REM sleep. Scientific Reports, 6(1), 1–13. https://doi.org/10.1038/SREP39229;TECHMETA=141;SUBJMETA=1595,2638,2649,2811,378,477,631;KWRD=COGNITIVE+NEUROSCIENCE,CONSOLIDATION,HUMAN+BEHAVIOUR

Oostenveld, R., Fries, P., Maris, E., & Schoffelen, J. M. (2011). FieldTrip: Open source software for advanced analysis of MEG, EEG, and invasive electrophysiological data. Computational Intelligence and Neuroscience. 10.1155/2011/156869

Rasch, B., Büchel, C., Gais, S., & Born, J. (2007). Odor cues during slow-wave sleep prompt declarative memory consolidation. Science (New York, N.Y.), 315(March), 1426–1429. 10.1126/science.1138581

Schreiner, T., Doeller, C. F., Jensen, O., Rasch, B., & Staudigl, T. (2018). Theta Phase-Coordinated Memory Reactivation Reoccurs in a Slow-Oscillatory Rhythm during NREM Sleep. Cell Reports, 25(2), 296–301. 10.1016/j.celrep.2018.09.037

Schreiner, T., Lehmann, M., & Rasch, B. (2015). Auditory feedback blocks memory benefits of cueing during sleep. Nature Communications. 10.1038/ncomms9729

Schreiner, T., Petzka, M., Staudigl, T., & Staresina, B. P. (2021). Endogenous memory reactivation during sleep in humans is clocked by slow oscillation-spindle complexes. Nature Communications. 10.1038/s41467-021-23520-2

Schreiner, T., Petzka, M., Staudigl, T., & Staresina, B. P. (2023). Respiration modulates sleep oscillations and memory reactivation in humans. Nature Communications, 14(1), 1–11. https://doi.org/110.1038/S41467-023-43450-5;TECHMETA=26,9;SUBJMETA=1385,1595,2638,2649,378,631;KWRD=CIRCADIAN+RHYTHMS+AND+SLEEP,COGNITIVE+NEUROSCIENCE,CONSOLIDATION

Schwartz, S., Clerget, A., & Perogamvros, L. (2022). Enhancing imagery rehearsal therapy for nightmares with targeted memory reactivation. Current Biology : CB, 32(22), 4808-4816.e4. 10.1016/J.CUB.2022.09.032

Shanahan, L. K., Gjorgieva, E., Paller, K. A., Kahnt, T., & Gottfried, J. A. (2018). Odor-evoked category reactivation in human ventromedial prefrontal cortex during sleep promotes memory consolidation. ELife, 7, 1–21. 10.7554/eLife.39681

Shen, X., Tao, L., Chen, X., Song, S., Liu, Q., & Zhang, D. (2024). Contrastive learning of shared spatiotemporal EEG representations across individuals for naturalistic neuroscience. NeuroImage, 301, 120890. 10.1016/J.NEUROIMAGE.2024.120890

Shi, X. S., She, Q., Fang, F., Meng, M., Tan, T., & Zhang, Y. (2024). Enhancing cross-subject EEG emotion recognition through multi-source manifold metric transfer learning. Computers in Biology and Medicine, 174, 108445. 10.1016/J.COMPBIOMED.2024.108445

Staresina, B. P., Bergmann, T. O., Bonnefond, M., Van Der Meij, R., Jensen, O., Deuker, L., Elger, C. E., Axmacher, N., & Fell, J. (2015). Hierarchical nesting of slow oscillations, spindles and ripples in the human hippocampus during sleep. Nature Neuroscience. 10.1038/nn.4119

van der Heijden, A. C., van der Werf, Y. D., van den Heuvel, O. A., Talamini, L. M., & van Marle, H. J. F. (2024). Targeted memory reactivation to augment treatment in post-traumatic stress disorder. Current Biology, 34(16), 3735-3746.e5. 10.1016/J.CUB.2024.07.019

Vaswani, A., Shazeer, N., Parmar, N., Uszkoreit, J., Jones, L., Gomez, A. N., Kaiser, Ł., & Polosukhin, I. (2017). Attention Is All You Need. Advances in Neural Information Processing Systems, 2017-December, 5999–6009. https://arxiv.org/pdf/1706.03762

Walker, M. P., & van der Helm, E. (2009). Overnight Therapy? The Role of Sleep in Emotional Brain Processing. Psychological Bulletin. 10.1037/a0016570

Wassing, R., Lakbila-Kamal, O., Ramautar, J. R., Stoffers, D., Schalkwijk, F., & Van Someren, E. J. W. (2019). Restless REM Sleep Impedes Overnight Amygdala Adaptation. Current Biology, 29(14), 2351-2358.e4. 10.1016/j.cub.2019.06.034

Wiesner, C. D., Pulst, J., Krause, F., Elsner, M., Baving, L., Pedersen, A., Prehn-Kristensen, A., & Göder, R. (2015). The effect of selective REM-sleep deprivation on the consolidation and affective evaluation of emotional memories. Neurobiology of Learning and Memory, 122, 131–141. 10.1016/j.nlm.2015.02.008

Xia, T., Chen, D., Zeng, S., Yao, Z., Liu, J., Qin, S., Paller, K. A., Platas, S. G. T., Antony, J. W., & Hu, X. (2024). Aversive memories can be weakened during human sleep via the reactivation of positive interfering memories. Proceedings of the National Academy of Sciences of the United States of America, 121(31). 10.1073/PNAS.2400678121

Xia, T., Yao, Z., Guo, X., Liu, J., Chen, D., Liu, Q., Paller, K. A., & Hu, X. (2023). Updating memories of unwanted emotions during human sleep. Current Biology, 33(2), 309-320.e5. 10.1016/J.CUB.2022.12.004

